# IMO-HIP 2015 Report: An Evolutionary Game Theory Approach to evolutionary-enlightened application of chemotherapy in bone metastatic prostate cancer

**DOI:** 10.1101/030262

**Authors:** Pranav Warman, Arturo Araujo, Conor Lynch, David Basanta

## Abstract

Prostate cancer metastasis to the bone is predominantly lethal and results from the ability of successful metastatic prostate cancer cells to co-opt microenvironmental cells and processes involved in bone remodelling. Understanding how the interactions between tumour and stromal cells determine successful metastases and how metastatic tumours respond to treatment is an emergent process that is hard to asses biologically and thus can benefit from mathematical models. In this work we describe a mathematical model of bone remodelling and the establishment of a prostate cancer metastasis in the bone using evolutionary game theory. We have mathematically recapitulated the current paradigm of a vicious cycle driving the tumor growth and we have used this tool to investigate the key interactions between the tumour and the bone stroma. Crucially, the model sheds light on the role that the interactions of heterogeneous tumor cells with the bone microenvironment have in the treatment of cancer. Our results show that resistant populations naturally become dominant in the metastases under a number treatment schemes and that schedules designed by an evolutionary game theory approach could be used to better control the tumour and the associated bone growth than the current standard of care.

## 1 Introduction

Prostate cancer is one of the most common types of cancer, with over 220,000 men diagnosed annually in the US alone, of which over 27,000 will succumb to it [1].

Virtually every patient that succumbs to prostate cancer does it as a consequence of metastatic growths in other organs. Furthermore, over 90 % of patients that die of prostate cancer show evidence of metastases to the bone. For this reason, a better understanding of how prostate cancer cells successfully metastasis to the bone is key if we are to improve how bone metastatic prostate cancer (mPCa) patients are treated. Improvements in our understanding of the molecular mechanisms involved have lead to the discovery of new putative therapeutical targets. But understanding the impact of treatments in a complex heterogeneous tumour requires new models that can incorporate several scales of biological insights to recapitulate the emergent process of cancer progression. In the past we demonstrated how a sophisticated but complex agent-based computational model could help us understand the role of tumor-host interactions [2]. While sophisticated agent-based models can accurately and quantitatively recapitulate cell dynamics in a specific area of a tumour, capturing the relevant intra-tumour heterogeneity and sensitivity to treatments can be a challenging and time-consuming process. Furthermore, simpler models can be used to represent the entire metastatic burden of the disease and thus could be used as proxies for the patient in optimisation algorithms [3]. For these reasons simpler qualitative models can be useful in providing an understanding of how certain interactions can shape evolution and resistance in cancer.

Previously Ryser and colleagues have used simple (ODE-based) mathematical models to understand bone remodelling. In this paper we use a mathematical model based on evolutionary game theory to investigate how prostate cancer tumour cells co-opt cells from the Basic Multicellular Unit (BMU), an area of trabecular bone that is being remodeled, to promote their own growth. The model recapitulates bone homeostasis in the absence of tumor cells. Bone homeostasis and its disruption by cancer has already been investigated by Dingli and colleagues [4]. But for the first time in a mathematical model of cancer based on game theory, we explicitly consider the role of the physical microenvironment in the progression of the tumour. Our model shows that tumor cells can interrupt the process of bone remodeling and co-opt the bone resident cells, the Osteoblast (OB) and Osteoclast (OC) into a vicious cycle of bone remodeling, where bone, a common good, is continuously being manipulated into being excessively created. Our results suggest that the pressure exerted by standard of care treatments, such as continuous chemotherapy, can lead to the rapid selection of resistant tumor cells. We propose that by discretising and modulating the treatment, it is possible to have a better control over the tumor growth.

### 1.1 Evolutionary Game Theory

Game Theory was initially developed by John Von Neumann and Oskar Morgernstern [5] to model problems in the fields of sociology and economics. Evolutionary Game Theory (EGT) is a branch of the game theory developed by John Maynard Smith [6] to better understand evolutionary dynamics in biological populations. Over the last decade and a half, research has emerged to use EGT to understand somatic evolution in cancer [7, 8, 9]. The main focus of these EGT models of cancer is how the interactions between different cell types in a heterogeneous tumour explain progression towards more aggressive cancers as well as towards treatment resistance. This focus on interactions with a heterogeneous population has recently gained increased focus thanks to the discovery of strong clinical evidence to suggest the importance of heterogeneity and Darwinian evolution in cancer [10].

## 2 The Model

### 2.1 The Environment

Our model aims to characterise homeostasis disruption by prostate cancer cells so the first step is to describe how this bone homeostasis is modelled. Bone remodelling is one of the processes that make the bone a very dynamic organ [11]. This process, involves a number of cellular species that interact together through a number of signalling molecules to orchestrate bone remodelling. It is known that this complex environment allows tumour cells to come in and co-opt the process for their own benefit [12]. We assume the tumor exists in the bone remodeling process (BMU) which is dominated by the stromal cells, specifically Osteoclasts and Osteoblasts, which remove and create bone, respectively. It is known that their is strong interplay between these two cells with the Osteoclasts releasing growth factors for Osteoblast generation and Osteoblasts release RANK-L which promotes Osteoclast differentiation and survival. In order to better characterise the interactions and proportions of the Osteoblasts and Osteoclasts we define functions that correlate their fitness to the amount of bone present:

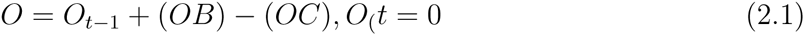

Additionally, as the variables OB and OC represent proportions, it is important to be able to discern a large proportion and small number of cells and large proportion and large number of cells as they would have quite different effects on the change of bone. Thus, we introduce coefficients to the above equation to weight the proportions with their prospective growth, the relative fitness.

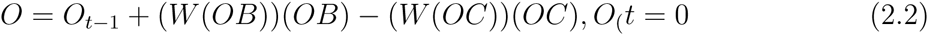

The functions that describes the bone-derived benefit or cost for OBs and OCs are symmetrical. Net bone formation is a function of the relative proportion of OBs and OCs in the population. The fitness of OCs and OBs are then inversely correlated so that the more bone the higher the fitness of OCs and the lower the fitness of OBs. This intuitively leads to a robust homeostasis. So bone-derived benefit for OCs could be described as:

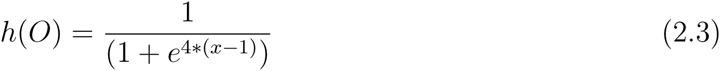

Respectively, the bone-derived fitness of OBs can be described as follows:

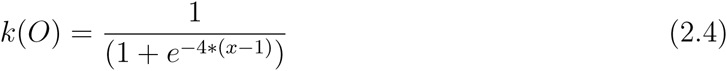

Thus, the replicator equations for OBs and OCs will be based on these relative fitness:

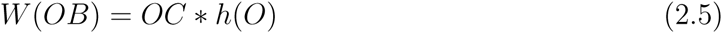

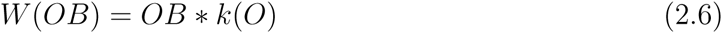

From these equations we can then generate plots showing the change in phenotype proportions over time. Given the way we have defined bone formation we can also plot net bone change over time.

From the replicator equations we can calculate the proportions of the next timestep by the following:

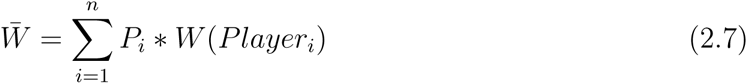

Where P is the proportion of Player*_i_*. And:

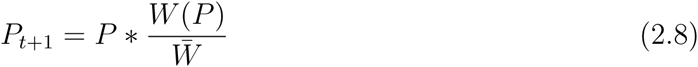

Thus for the replicator equations 2.6 and 2.7 we get the plots shown in figure 1.

**Figure 1:**
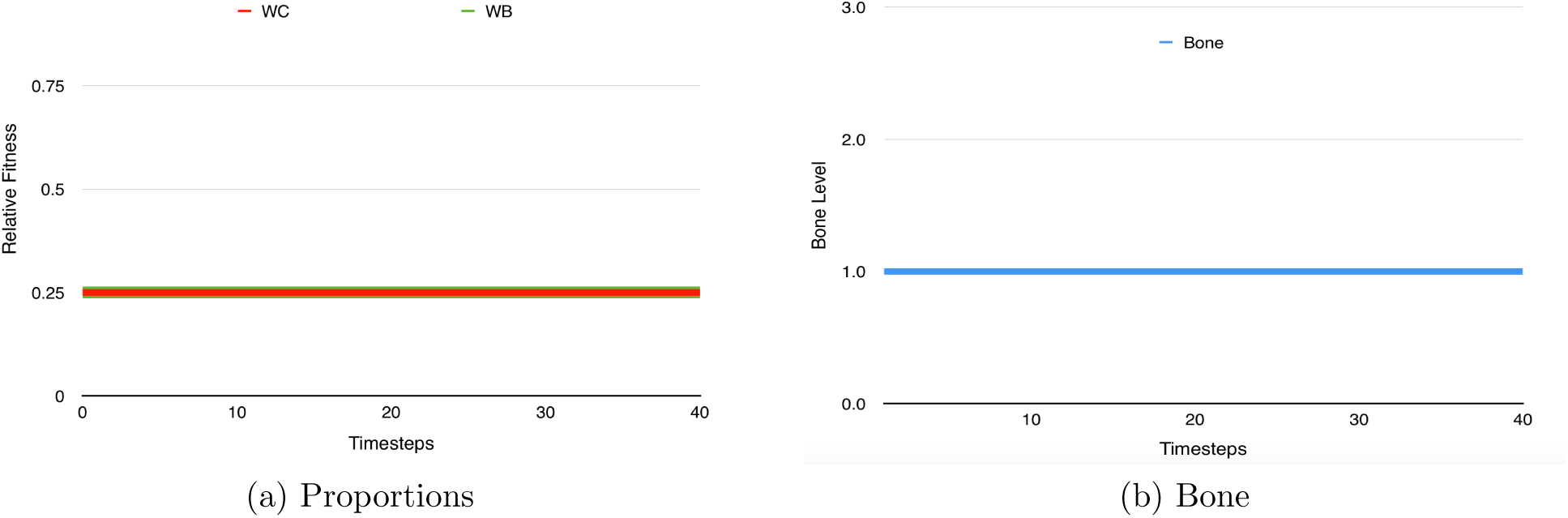
Bone homeostasis. The Osteoblasts and Osteoclasts quickly form equal proportions and in the event of a disruption attempt to return to this normalcy. Initialization values: *P_OB_* = *P_OC_* = 0.

### 2.2 Tumor Introduction

Now that we have established a working model for bone homeostasis, we can begin to investigate how the game changes when metastatic prostate cancer cells are introduced:

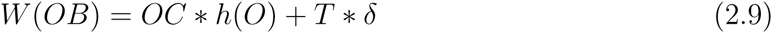

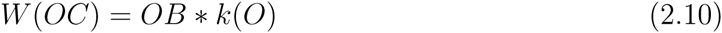

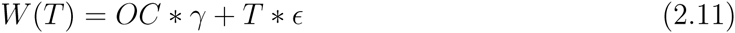

We define γ to be the benefit the tumor receives from the OCs breaking the bone to release growth factors including TGF-*β, δ* to be the benefit OBs receive as a result of their interactions with tumor cells mediated by signalling molecules such as TGF-*β* and RANKL [Bussard], and *ϵ* as the benefit the tumor receives the growth factors released by other tumor cells. γ ≫ δ ≫ *ϵ*. We use these equations to generate the plots seen in figure 2.

**Figure 2:**
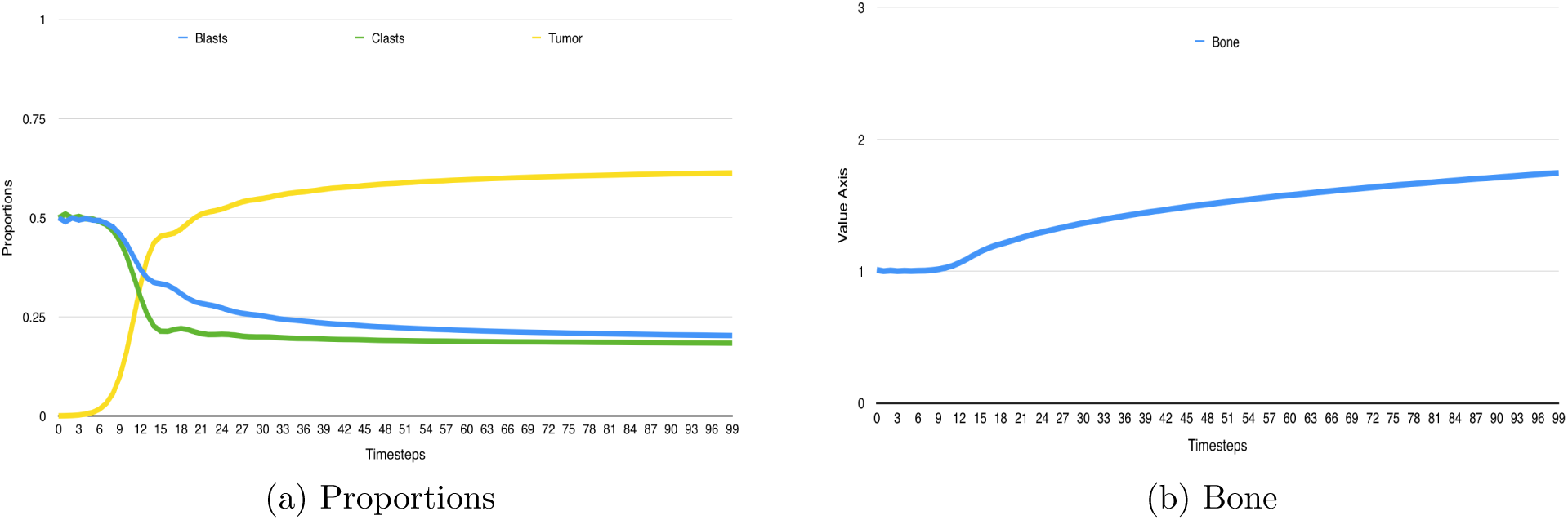
Homeostatic disruption due to tumour introduction. The simulation begins primarily with Ostebolasts and Osteoclasts with a very small proportion of tumor cells, which due to a higher relative fitness dominates and causes the bone to surge. Initial values: *P_OB_* = *P_OC_* = 0.4998; *P_T_* = 0.0004; γ = 0.95; ϵ = 0.03; *δ* = 0.2.

## 3 Chemotherapy

Chemotherapy is part of the standard of care for metastatic prostate cancer patients and is the main tumor treatment we will consider in the remaining of this report.

### 3.1 Model

Chemotherapy targets cells which are proliferating, and the quicker the proliferation, the more that phenotype is attacked by the treatment. Thus in order to apply chemotherapy to our model takes into account the impact of the treatment not only the tumor population, but also on the Osteoclasts and Osteoblasts. This is done by reducing the relative fitness for the affected phenotypes by multiplying an equivalent cost to the replicator equations as such, the larger the cost, the more efficacious the treatment scheme. This equivalent cost has a larger effect on the tumor population due to its larger relative fitness - a proxy for the player’s proliferation rate.

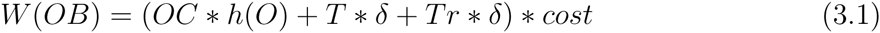

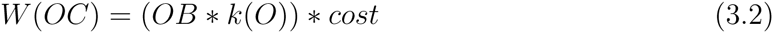

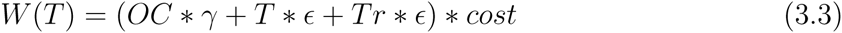

These equations can be used to produce figure 3. Though the treatment is initiated we see no effect on the tumour proportion, and we are only cued to a change by the substantial drop in the players’ relative finesses. The unchanging proportion and falling of the relative fitnesses forces us to think of the behavior of the count of cells - falling - as opposed to the presented proportion of cells - unchanged.

**Figure 3:**
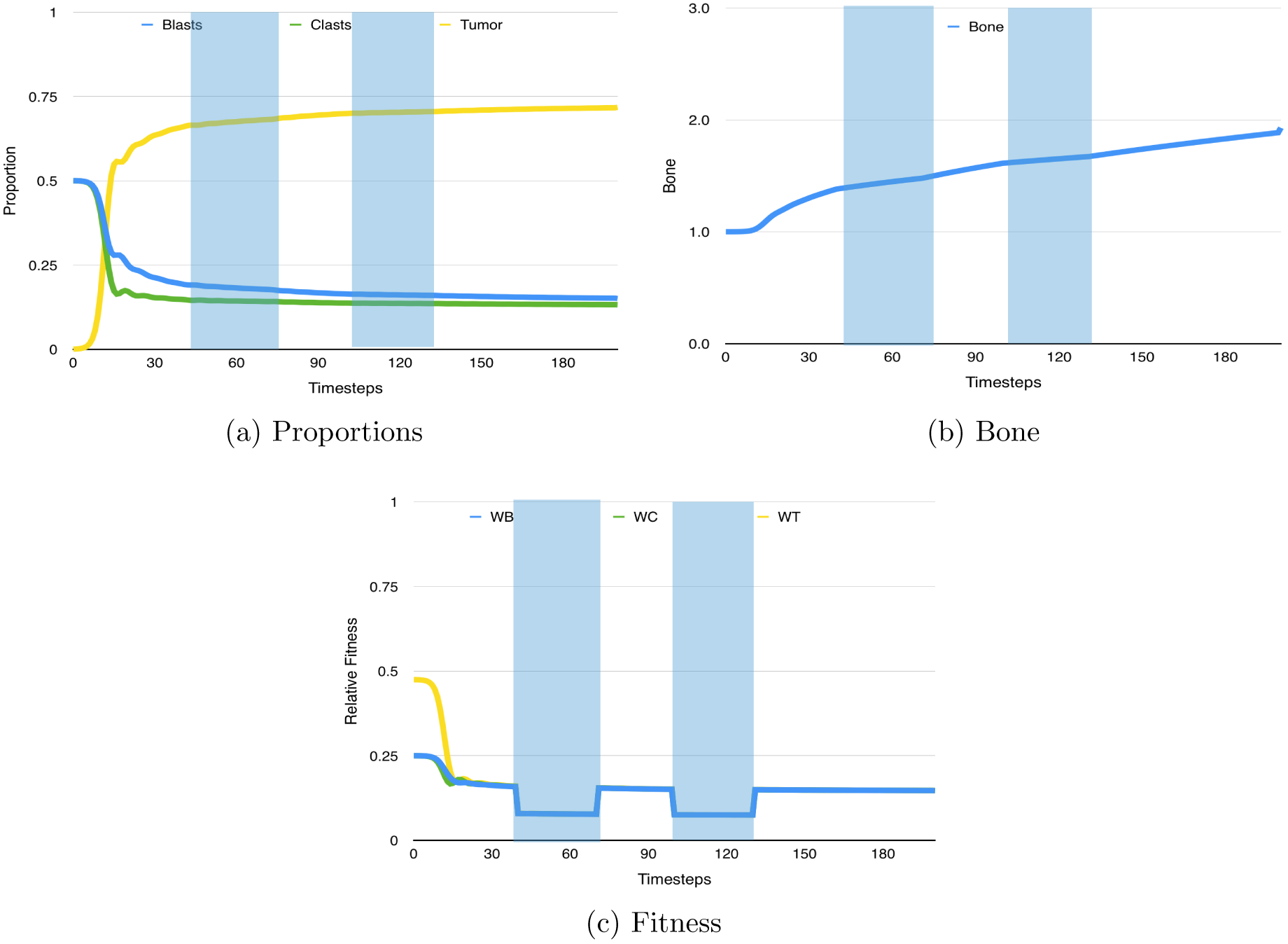
Chemotherapy on a homogeneous tumour population. We see that the treatment has no effect on the proportions of the cells, but it effects the amount as shown by the drop in the player’s relative fitness. Initial values: *P_OB_* = *P_OC_* = 0.4998; *P_T_* = 0.0004; γ = 0.95; ϵ = 0.03; *δ* = 0.2. Treatment began at timestep 40, for two 40 timestep treatments with a 20 timestep interval and an efficacy of 0.5.

### 3.2 Resistant Tumor Population

However, it is also well observed and accepted that during treatment schemes the composition of the tumour changes as parts of the tumour population becomes immune to the treatment, often due to a mutation. Thus, in order to improve the reality of our model we include this population, changing the equations to:

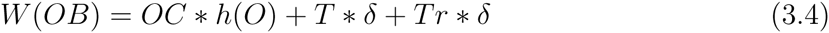

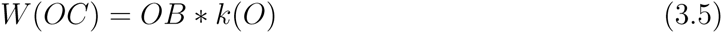

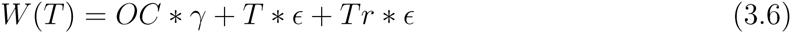

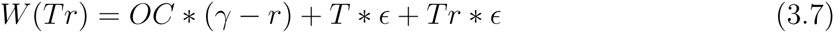

The variable for *γ*, *δ*, *ϵ* are kept the same as the susceptible tumor population as both tumor populations exhibit similar lifestyles; however the tumor’s which are resistant (Tr), are so due to a different pathology which while having a benefit during treatment schemes, extricate a cost from the cell, which lowers the cell’s relative fitness. This is shown in the replicator equations above through the subtraction of a cost ”r” from the *γ* to lower the Tr’s fitness. These equations are then used to create plots as shown in figure 4.

**Figure 4:**
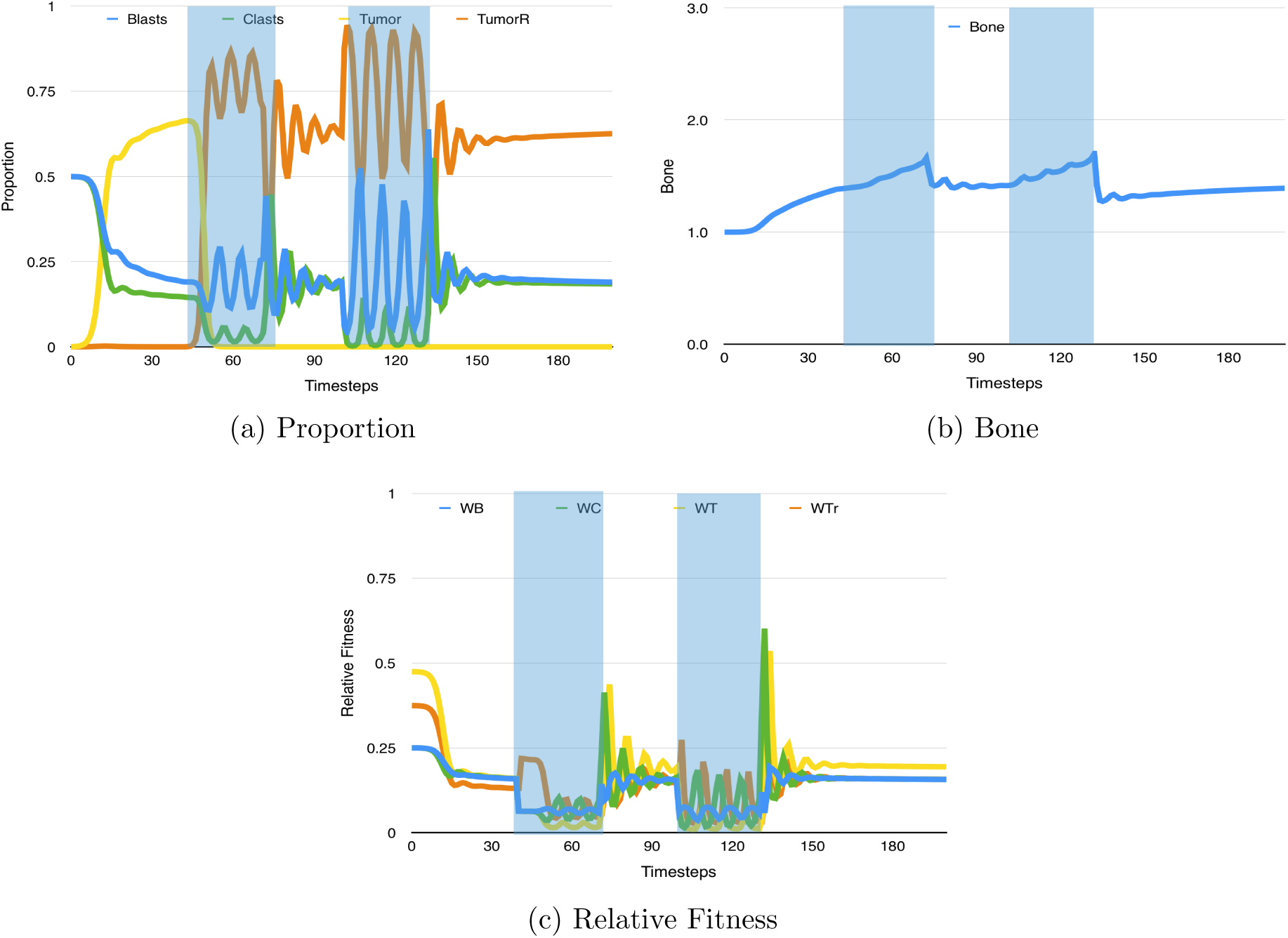
Chemotherapy on a heterogeneous population. The bone level begins to temporarily decrease after treatment due to decreased level of tumour and increased level of stromal cells.Initial values: *P_OB_ = P_OC_* = 0.4998; *P_T_* = 0.0004; *r* = 0.1; γ = 0.95; *e* = 0.03; *δ* = 0.2. Treatment began at timestep 40, for two 40 timestep treatments with a 20 timestep interval and an efficacy of 0.5.

In fig. 4 we see the change in the composition of the tumour as the resistant population takes over. In some cases, as seen in fig 4, the susceptible tumour population becomes dominant once again due to it’s small but present proportion and a larger relative fitness than other players.

### 3.3 Model Results

The starting point of our work was to ensure that the mathematical model would recapitulate bone homeostasis in the absence of interruptions by tumour cells. Fig. 1 shows the results of the replicator equation with only OBs and OCs. Although OBs and OCs do not interact directly, they maintain bone homeostasis over time and their proportions, although cyclical, remain similar over time. This homeostasis is disrupted by the inclusion of tumour cells. Figure 2 shows how the introduction of this population, and the assumption that T and OB populations work in a synergistic fashion, leads to a vicious cycle where tumour cells become the majority and where, given the relative abundance of OBs over OCs, bone keeps growing over time leading to growth in pathological bone.

The application of chemotherapy will impact the tumour but, as figure 3 shows, this might not change the relative proportions of the different populations. Since chemotherapy is assumed to impact rapidly proliferating cells equally (we do not assume that stromal cells are more resistant to chemotherapy than tumour cells, only that they usually proliferate at a lower rate), the impact of the treatment can be better seen in fig 3c. The success of the treatment can also be appreciated in fig. 3b as bone growth is halted during the treatment windows. A more realistic scenario is the one portrayed in fig. 4 where the tumour population is heterogeneous and made of tumour cells that are susceptible to chemotherapy (and thus pay a cost under the presence of treatment) and tumour cells that are chemoresistant (TR). Under this scenario, the application of chemotherapy changes the composition of the population by selecting for the TR phenotypes. If the application of chemotherapy is short enough, there might be enough susceptible tumour cells that could allow for this subpopulation to recover. Bone growth can be observed during, and in between, applications of chemotherapy.

All these examples highlight how a simple game theoretical model can be used to explore different schemes of chemotherapy application. These schemes were characterized by the variables of treatment interval, duration, efficacy and amount. We considered three types of treatment duration - short, average, and long. In each case the replicator equations were iterated 40 times before the treatment started with an initially small proportion of tumor cells (3.6 * 10^−4^) and a resistant tumor population of 4 * 10^−5^, in order to allow tumor growth as in reality treatments do not begin as soon as tumors appear but only after symptoms show up. We assumed a cost of resistance (r) to be 0.1 and *ϵ* to be 0.03. For each of these schemes 4 different values were recorded and grouped in figure 5: (1) susceptible tumour proportion, (2) bone growth, (3) average tumour fitness, and (4) average player fitness. The last two yield results that can be understood intuitively. As we increase drug efficacy we reduce the proportion of the population made of tumour cells (resistant and susceptible) and the amount of bone produced.

**Figure 5:**
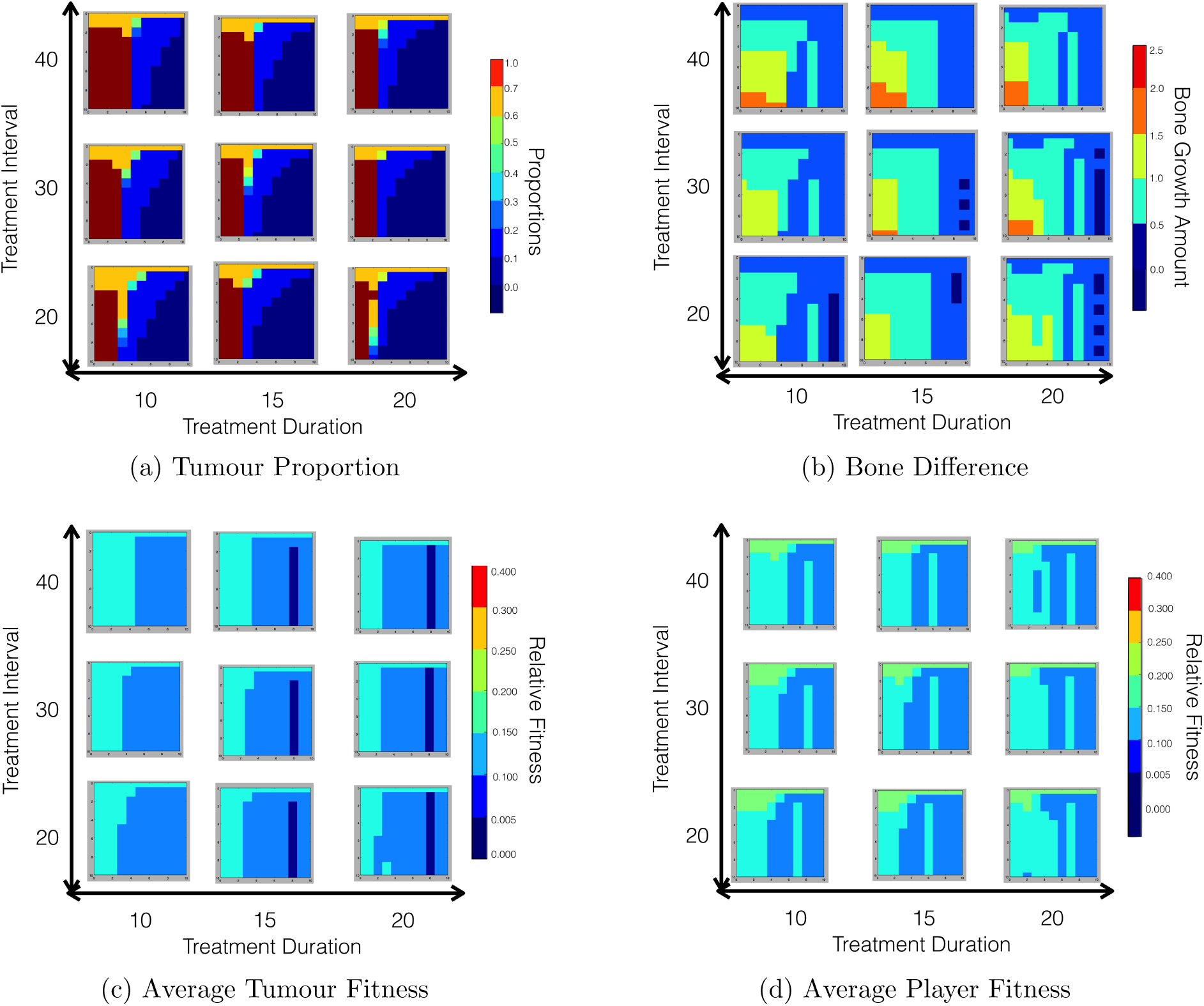
Chemotherapy on a heterogeneous tumour population. The (1) Tumor proportion, (2) difference in bone level after treatment, (3) average tumour fitness, and (4) average player fitness. We show the average fitness for the players in our model under different treatment schemes. The major horizontal axis on the outside represents the treatment duration and the major vertical axis on the outside represents the treatment interval. The smaller horizontal axis on each square is the treatment efficacy, incremented by 10 percent while the smaller vertical axis on each square is the amount of treatments given incremented from at the top 1 to 10. Initial values: *P_OB_ = P_OC_* = 0.4998; *P_T_* = 0.00036; *P_TR_* = 0.00004; *r* = 0.1; γ = 0.95; ϵ = 0.03; *δ* = 0.2.

### 3.4 Result Analysis

Regarding the tumour susceptible proportion: The values were taken exactly one timestep after the last treatment in order to best determine the outcome of the treatment. The general trend is for the tumour proportion to decrease as the treatment scheme becomes more intense (due to a higher efficacy and more treatments). The top bar is unaltered as it signals no treatment so despite an unrealistically high efficacy the tumour grows unhindered. Regarding the bone difference plot: The amount of bone growth shows a trend towards larger bone growth in light yet long treatments. During these schemes the susceptible tumour cells are given a longer duration of time, due to longer treatment schedules, and thus have longer perturbation on the OB fitness causing the rapid bone growth over a longer period of time. Additionally, with other long treatment schemes the treatment often eliminates the tumour susceptible and creates an ESS between the resistant phenotype and stromal cells such that bone re-normalizes and becomes static.

Additionally, we continue studying the aforedescribed treatment schemes through another perspective: the average fitness, which tells us more about the model and the potential and rate of growth and when combined with our data of the proportions gives us a better conception of how the players (stromal and tumor cells) are interacting. It is of no surprise when looking that both the average tumor fitness and the average player fitness behave in the similar fashion of decreasing as the treatment intensifies. The dark blue line that is seen in the average tumor fitness subplot is due to disproportionately high OB fitness in the disrupted cycle of OB and OC proportions which causes the tumor fitness to drop - however, in other cases the efficacy is either too high or too low to cause the cycle’s nadir to occur right at the end of the treatment.

## 4 Adaptive Therapy

Adaptive Therapy (AT) is an evolutionary enlightened treatment scheme in which the goal of the treatment is not to kill the tumour, but rather control its size. By controlling the tumour population we prevent the resistant population from becoming the dominant phenotype, while still controlling the overall tumour burden.

This treatment would be applied by checking the patients tumor population and giving treatment only when the tumour population grows over a certain size. In this EGT model we assume that the fitness of the tumour population constitutes a proxy to tumour size. However, this would involve checking the patients tumor levels on a daily basis which is not always possible. Thus, in order to apply this treatment we modify it in the following way:

1. Chemotherapy will start as in other treatment schemes at a late timestep such that the tumor proportion has achieved some form of evolutionary stable proportion, mirroring a fully grown tumor.
2. The patient’s tumor proportion will be checked three times a week and if the tumor proportion is higher than a set proportion the treatment, much like a boolean, will be turned off and will only be turned on if during another checkup the tumor proportion crosses the predefined value

Using this modified Adaptive Therapy treatment scheme, we are able to generate the results shown in figure 6.

**Figure 6:**
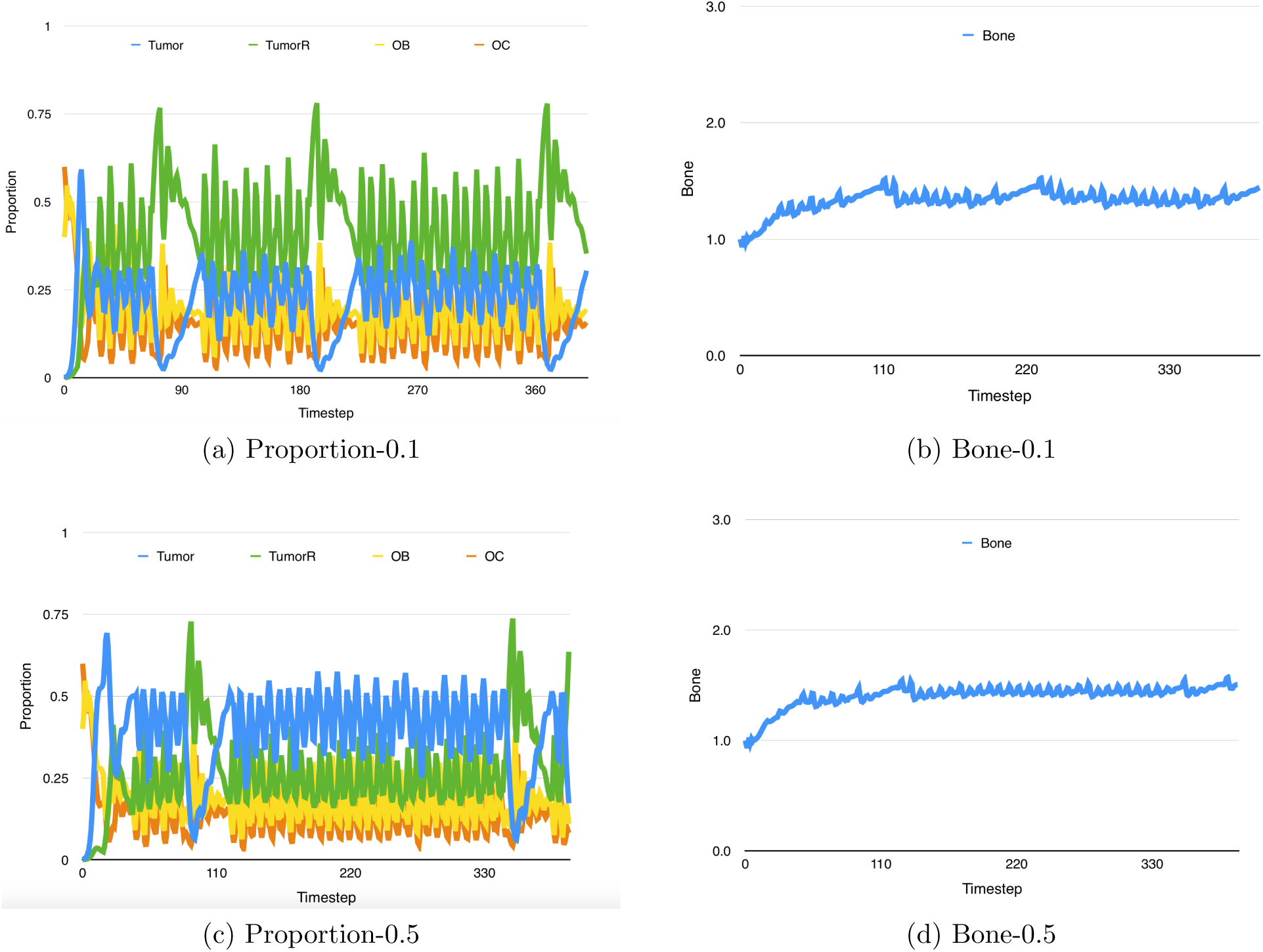
Adaptive Therapy controls the Tumor proportion to be less than 0.5. The growth of the heterogeneous tumor population remains checked. These plots also show the effect of such a treatment on the amount of bone. Initial values: *P_OB_ = P_OC_* = 0.4998; *P_T_* = 0.00036; *P_Tr_* = 0.00004; *r* = 0.1; γ = 0.95; ϵ = 0.03; *δ* = 0.2.

**Figure 7:**
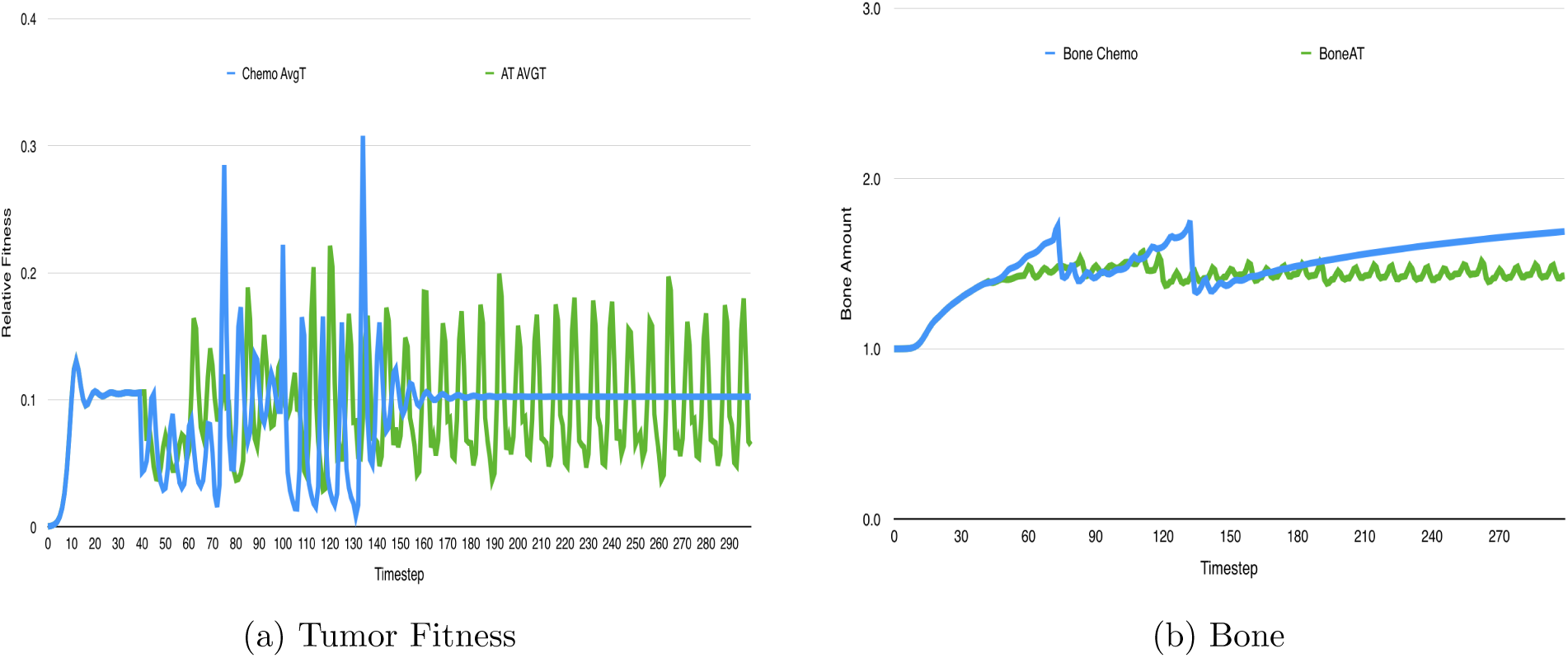
Tumor fitness for a classic treatment versus adaptive therapy. Initial values: *P_OB_ = P_OC_* = 0.4998; *P_T_* = 0.00036; *P_Tr_* = 0.00004; *r* = 0.1; γ = 0.95; ϵ = 0.03; *δ* = 0.2.

### 4.1 Model Results

We see that the realistic Adaptive Therapy method which we outline above is successful at controlling the bone growth and changes the cancer from a fatal disease to a chronic problem. Additionally, we observe that, based on what values are chosen for control, the tumor burden can be controlled and the resistant population can be controlled and prevented from completely displacing the susceptible population The full impact of the AT treatment scheme can be seen in figure 6, where the average tumor fitness for an Adaptive Therapy treatment (with the with the tumor controlled to 0.5) is overlayed on the average tumor fitness. Here we see that as soon as treatment is the average tumor fitness (avgT) falls followed by controlled oscillations of the AT avgT compared to oscillations and large spikes of the Chemotherapy avgT. Additionally, we show the bone growth over time for both of these treatment schemes, and as expected, the bone under the adaptive therapy is controlled, while the bone under standard chemotherapy increases indefinitely. Within the bone plot we note two spikes in the bone under the chemotherapy.

## 5 Discussion

The results presented in this report were collected during the HIP-IMO in June-August 2015. During these months we developed a qualitative mathematical model of bone remodeling that involves the interactions between bone-producing cells (osteoblasts) and bone-destroying cells (osteoclasts). While qualitatively modeling bone homeostasis with these two cell types is not novel [4], our approach explicitly incorporates, for the first time in a mathematical model of cancer based on game theory, the microenvironment. This is important because the model we presented allows us to study not only bone homeostasis but also homeostasis disruption as a result of metastasis from the bone and the impact of such metastasis in the bone. Our results recapitulate not only this (vicious cycle) but also the bone formation that characterizes prostate metastases in the bone [1].

The heterogeneity of tumours is increasingly recognized as an important driver of evolution of resistance in cancer. Chemotherapy is one of the most widely used treatment options in metastatic prostate cancer. Resistance to chemotherapy is common and results from the treatment providing strong selection for tumor cells that can avoid its cyto-toxic effects. This resistance is often obtained through the upregulation of drug exporter pumpts on the surface of tumour cells. Producing and maintaining these pumps is energetically costly to the tumour cells but allow those that have many of them to deal with many of the drugs commonly used in the treatment of metastatic prostate cancer. Our results show that conventional chemotherapy-based treatments that aim to reduce tumour burden as much as possible work well if the metastases are homogeneous. Assuming heterogeneity with regards to chemotherapy resistance, our model allows us to explore how different treatment durations and intervals between drug application impact tumour heterogeneity, fitness, and the increase in bone formation (see fig. 5).

While some of the results are very intuitive, some of them are not. The number of treatment intervals for instance, while having an impact on the tumour fitness, also impacts the formation of pathological bone. This is not obvious from the model assumptions and is a result of clinical significance as many treatments for metastatic prostate cancer patients are palliative [13] and pathological bone is a major factor impacting negatively patient’s quality of life.

While the standard of care involves the use of treatments like chemotherapy where dosage is determined by maximum tolerable dose, new treatment strategies have been recently proposed that could be described as evolutionary enlightened. One of them is known as adaptative therapy (AT) and the aim is to control the tumour population rather than to reduce the burden. AT have been explored mathematically by Gatenby and colleagues [14] but under the assumption that the microenvironment of the tumour is not relevant and that tumour populations do not interact with each other. Our investigation of AT on prostate cancer metastases to the bone assumes that we can track metastatic burden with PSA (Prostate Specific Antigen). Results (see fig. 6) suggest that AT can indefinitely control the tumour population (assuming that chemotherapy resistance is costly even if easy to evolve). Furthermore, we have also shown that compared to standard chemotherapy, AT can control bone formation.

We intend to use this simple model as a theoretical platform in which to investigate other treatments of the standard of care for metastatic prostate cancer patients. These include, but are not limited, to biphosphonates, anti-RANKL and hormonal therapy. These will allow us to understand how the combination of treatments in heterogeneous bone metastases would allow us to control metastatic burden and pathological bone growth.

## 6 Acknowledgments

This work was performed by Pranav Warman under the supervision of David Basanta and in collaboration with Arturo Araujo and Conor Lynch. PW and DB wish to thank HIP (High school Internship Program) and its director, Heiko Enderling, for making this work possible. They wish to thank the Integrated Mathematical Oncology department for hosting and facilitating HIP and the Moffitt Cancer Center for generously funding it.

